# Regulation Flow Analysis discovers molecular mechanisms of action from large knowledge databases

**DOI:** 10.1101/2025.06.15.659785

**Authors:** Carlos P. Roca, Oleg Sysoev, Elena Eyre, Silvia Galan, Dominic Sinibaldi, Philip Tedder, Jonathan Mangion

## Abstract

Drug development is a long and expensive process, with only a small fraction of potential drugs being finally approved. The challenge of drug development is rooted in our limited understanding of biological systems and the disease processes that drugs are trying to modulate. We propose a novel method, called Regulation Flow Analysis (RFA), which is based on the principles of biological regulation, causal graphs, and graph flow. RFA is able to generate causal hypotheses of mechanism of action, using large Knowledge Graphs (KG) of molecular regulation. Our numerical experiments demonstrate that the generated hypotheses, in the form of regulation pathways, often summarize mechanisms of drug action in a manner both understandable and actionable. Thus, RFA can greatly improve our understanding of the biological processes underlying health and disease, and therefore substantially facilitate drug development. Our examples illustrate how RFA recovers known mechanisms of action and can be used for target and biomarker discovery and validation.

## 1 Introduction

The process of drug discovery and development is both long and expensive, with an average time of 15 years for a drug to reach market, at a cost of approximately two billion dollars [1]. Less than 10% of new drugs in Phase 1 will eventually be launched, with the majority of failures happening in the transition from Phase 2 to Phase 3, because of toxicity or lack of efficacy [2, 3]. The difficulty of drug development is linked to the inherent complexity and incomplete understanding of the biological systems that are being targeted. If reliable hypotheses of the mechanisms of action of potential drug targets can be developed and tested, this will lead to higher success rates due to better target discovery and validation. Additionally, it would lead to the selection of more accurate biomarkers and the better prediction of possible side effects.

AstraZeneca applies the 5R framework to R&D [4]: right target, right tissue, right safety, right patient, and right commercial potential. Target selection remains the most important decision in the drug discovery process, and interrogating the growing knowledge of disease biology is critical to making the right choice. Information and knowledge about diseases, targets and drugs is available from many different sources, but integrating this biomedical knowledge is complex and challenging. Here we focus on the critical aspect of improving our understanding of the underlying disease processes and the ways to modulate them, by generating accurate pathways between genes or proteins, as causal graphs representing biological regulation. This allows the rational selection of drug targets, the prediction of downstream events of modulating those targets, and the identification of potential biomarkers.

Regulation networks can be represented by directional and causal graphs, which provide the proper conceptual framework to model known interactions between biological entities, such as protein-protein interactions, post translational modifications, and transcriptional regulation. These forms of knowledge graphs (KGs) can be readily built from available data resources, both academic (for example, OmniPath [5] or Reactome [6]) and commercial (such as Clarivate MetaBase [7], Qiagen Knowledge Base [8], or Metaphacts Knowledge Graph [9]). These knowledge bases collect large amounts of information in the literature and in curated databases about biological regulation at molecular level. Being very information rich, such large KGs have a great potential for revealing the mechanisms of action of potential drug targets.

Signaling in large KGs has been modeled by a number of approaches that integrate omics data [10, 11, 12, 13, 14, 15, 16]. Ingenuity Pathway Analysis (IPA) [17] computes upstream regulators of the given gene set and generates mechanistic hypotheses connecting the upstream regulators with this gene set. Similar principles are also used for identification of the downstream targets in IPA. When it comes to the methods that generate mechanistic hypotheses purely from KGs, the literature is very scarce. Nichenet applies Dijkstra’s shortest path algorithm to infer signaling paths between a ligand and the target gene of interest [14]. Signal Flow Control (SFC) algorithm [18, 19] introduces a dynamic model with a combination of network signal propagation and basal activation, and it uses the gradients of the network flow to estimate the signal in the steady state. This method demonstrates that the sign of the predicted effect of a given perturbation agrees with the direction of the experimentally observed differential expression changes in 60-80% of genes. While the objective of SFC is not the generation of mechanistic hypotheses, it illustrates that algorithms on regulation graphs are able to explain the effects observed experimentally to a reasonable extent.

In this paper, we present a novel method, called Regulation Flow Analysis (RFA), developed and actively used in AstraZeneca for generating hypotheses of mechanism of action at molecular level, by identifying the causal events connecting sources of perturbation (i.e. a deleterious genetic variant, or a ligand-receptor binding) to the observed effects (a change in gene/protein expression, or a change in a molecular biomarker). Our model consists of two parts: the Regulation Flow model and the algorithm for generating hypotheses of mechanism of action. While previous work has explored the potential of graph flow to model biological regulation [18, 19], we consider the algorithm to generate mechanistic hypotheses as the main novel contribution reported in this paper. We apply our method to predicting the underlying biology of Noonan syndrome, thus demonstrating the capacity of RFA for generating mechanistic hypotheses, and to identifying drug targets for a developmental disease, Fragile X syndrome.

## 2 Regulation Flow model

To model regulation in KGs, we propose a model which we call Regulation Flow model. The model assumes that after one or a few vertices in the KG are perturbed, a cascade of regulation effects propagates throughout the graph. The model does not have a temporal meaning, but rather it represents the steady state some time after the perturbation happened. The effect on any given vertex will be, by assumption, the addition of the effects arriving by all the possible different paths in the graph connecting the initially perturbed vertices to that particular vertex.

In mathematical terms, we have a regulation graph *G* =*< V, E >*, where *V* = {*v*_1_, …, *v*_*n*_} is the set of graph vertices or nodes, and *E* = {*e*_*ij*_ = (*v*_*i*_, *v*_*j*_); *v*_*i*_, *v*_*j*_ ∈ *V*}, is the set of graph edges or links, with associated labels *l*_*ij*_. In general, vertices represent genes or proteins, whereas edges represent direct regulation relationships, at molecular level, between them. When *l*_*ij*_ = 1, we have positive regulation, this meaning that up-regulation of *v*_*i*_ leads to the up-regulation of *v*_*j*_, and conversely down-regulation of *v*_*i*_ leads to the down-regulation of *v*_*j*_. On the other hand, *l*_*ij*_ = −1 indicates negative regulation, so that if *v*_*i*_ is upregulated, then *v*_*j*_ is down-regulated, and conversely down-regulation of *v*_*i*_ causes up-regulation of *v*_*j*_. Up- and down-regulation have here the usual meaning in biological systems, that is, stimulation and inhibition, respectively.

Given these inputs, we compute a signed adjacency matrix *A* = (*A*_*ij*_) from the graph *G*, following the standard definition, *A*_*ij*_ = *l*_*ij*_ if (*v*_*i*_, *v*_*j*_) ∈ *E* and *A*_*ij*_ = 0 otherwise. By using the adjacency matrix, we compute an *unscaled normalized step matrix U* = (*U*_*ij*_) as the transpose of the *A* matrix, normalized by the number of outgoing edges of each vertex:

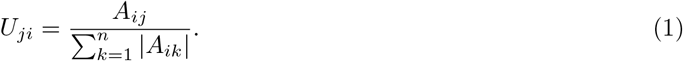

The rationale of this normalization is that vertices (genes or proteins) with effects on many others vertices, will tend to have effects of smaller magnitude or strength, in comparison to vertices affecting a lower number of vertices.

This unscaled step matrix represents the propagation of a regulation effect in a single step, or in other words, over a path of length one in the graph. In an experiment or intervention, we are given a perturbation set with one or a few vertices, 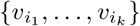, describing the vertices that are the origin or source of the perturbation. We codify this perturbation with the vector *p* = (*p*_*t*_), with *p*_*t*_ ≠ 0 when *t* ∈ {*i*_1_, …, *i*_*k*_} and usual values |*p*_*t*_| = 1. For example, a single perturbation may be a drug which stimulates a receptor, which in another intervention may be combined with a compound that inhibits a kinase. Let *x* = (*x*_1_, …, *x*_*n*_) represent the resulting regulation effect, so that *x*_*i*_ *>* 0 if the perturbation *p* up-regulates vertex *v*_*i*_, *x*_*i*_ *<* 0 if it down-regulates vertex *v*_*i*_, and *x*_*i*_ = 0 otherwise. Then, the one-step propagation of regulation can be calculated as:

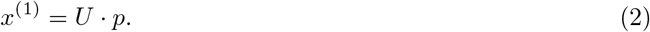

Let us also define *x*^(0)^ = *p*, the perturbation itself, without effects on any other vertex.

The regulation reaching a vertex after traversing *m* steps in the graph can also be calculated as:

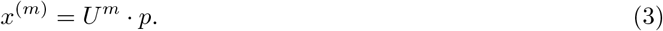

The *total regulation* on any vertex, composed by the regulation reaching via paths of any length, is accordingly:

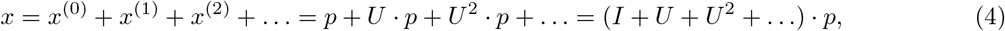

*I* being the identity matrix.

Since (4) includes the sum of an infinite series, convergence criteria must be verified. The sum of the matrix powers *I* + *A* + *A*^2^ + … is a Neumann series ([20], page 618), whose necessary and sufficient criterion of convergence is that the spectral radius *ρ*(*U*) (the modulus of the largest eigenvalue) is strictly less than one. This can not be guaranteed for matrix *U* in general, and therefore we introduce the *scaled step matrix S* as follows:

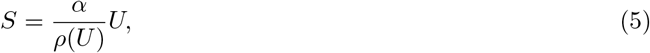

where 0 *< α <* 1, for example, *α* = 0.99. As by definition *ρ*(*S*) = *α*, this guarantees *ρ*(*S*) *<* 1. Accordingly, we modify equation (4) as follows:

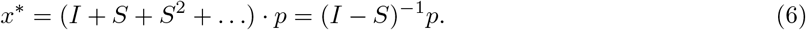

This way, the parameter *α* represents the scale of the one-step effect produced by perturbing with the largest eigenvector of the step matrices *U* or *S*. Therefore, it controls the overall scale of the response to perturbations in the model.

The last part of equation (6), as well as the existence of the inverse, follows from the properties of the Neumann series ([20], page 618). We call the matrix *F* the *regulation flow matrix*,

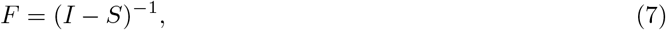

and *x*^∗^, the regulation effect caused by perturbation *p*, corresponds to the *total regulation flow*,

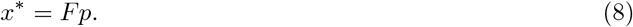

Each coefficient of the regulation flow matrix, *F*_*BA*_, represents the regulation flow from A to B, that is, the total regulation influence of gene/protein A over gene/protein B. A compact description of the Regulation Flow model is provided in Algorithm 1.

### Algorithm 1

Regulation flow model

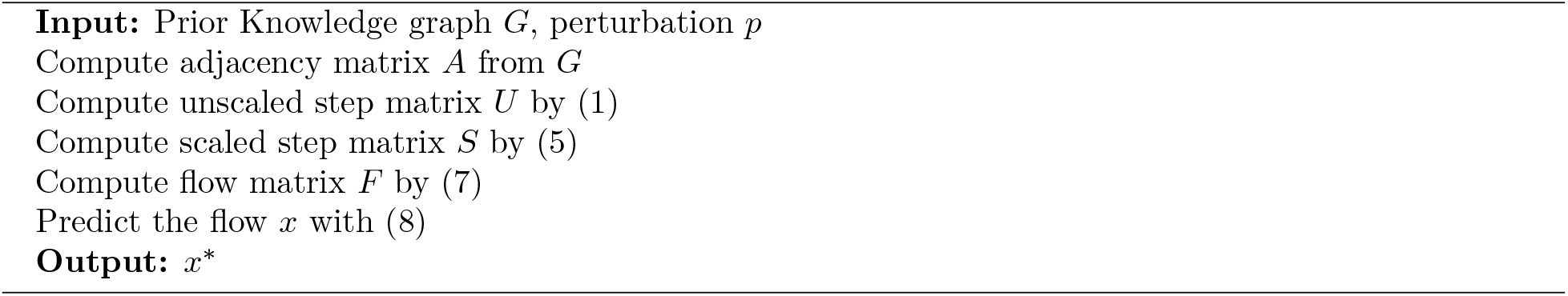

## 3 Generation of mechanistic hypotheses by RFA

Pathways are essential constructs in biology and biomedicine, conceived to organize biological processes as series of events describing the causal links between an initial perturbation and a series of downstream effects. By perturbation we mean, for example, the activation of a kinase, the coupling of a ligand with its receptor, or the formation of a protein complex. Here, we formalize the concept of a pathway with a precise definition. As a first step, we will define the *pathway from gene/protein A to gene/protein B* as the subgraph defined by all the regulation paths connecting *A* to *B*. In most practical applications such a pathway would contain a very large number of intermediate genes/proteins and regulation paths, the latter frequently infinite because of the presence of cycles. However, as we will show, a crucial property is that in most cases only a very small number of intermediate genes/proteins and regulation paths channel almost all of the regulation flow from *A* to *B*. This allows to consider the much simpler *effective pathways* between *A* and *B*, instead of the full pathway.

Let us start considering the intermediate genes/proteins {*C*_1_, *C*_2_, …, *C*_*n*_} between the genes/proteins *A* and *B*. For any intermediate gene/protein *C*_*i*_, there exists at least one path connecting *A* to *C*_*i*_, and at least one path connecting *C*_*i*_ to *B*. Any path *P* from *A* to *B* traversing through *C*_*i*_ will be defined by the sequence of vertices *P* = (*A, D*_1_, *D*_2_, …, *D*_*n*_, *B*), with at least one *j* = 1 … *n* such that *D*_*j*_ = *C*_*i*_. We call this path *C*_*i*_-acyclic if *C*_*i*_ occurs only once in the sequence *P*. In general, let us consider *k*_1_ to be the index of the first occurrence of vertex *C*_*i*_ in *P*, and let *k*_2_ be the index of the last occurrence of *C*_*i*_ in *P*. If *k*_1_ = *k*_2_, then *P* is *C*_*i*_-acyclic. Otherwise, when *k*_1_ ≠ *k*_2_, *P* is a concatenation of 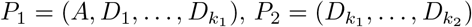 and 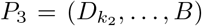, where *P*_1_ and *P*_3_ are *C*_*i*_-acyclic and *P*_2_ is a cycle from *C*_*i*_ to *C*_*i*_, which we call a *C*_*i*_-cycle. *C*_*i*_-cycles may have any finite number of visits to vertex *C*_*i*_.

From this, it is clear that all the paths from *A* to *B* that traverse through *C*_*i*_ can be obtained as the composition of all the *C*_*i*_-acyclic paths from *A* to *C*_*i*_, all the *C*_*i*_-cycles, and all the *C*_*i*_-acyclic paths from *C*_*i*_ to *B*. By making *C*_*i*_ = *A* or *C*_*i*_ = *B*, we can also state that all the paths from *A* to *B* can be obtained as the composition of all the *A*-cycles and all the *A*-acyclic paths from *A* to *B*, or as the composition of all the *B*-acyclic paths from *A* to *B* and all the *B*-cycles.

Algebraically, the total regulation flow from *A* to *B* is equal to the sum of the flow contributed by each of the connecting paths (4). Additionally, the flow channeled by any path *P* can be factored into the multiplication of the flow channeled by any two subpaths *P*_1_ and *P*_2_ such that *P* is the concatenation of *P*_1_ and *P*_2_. As a result, by the distributive property, the composition of paths corresponds algebraically to the multiplication of flow.

Let us denote by 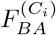 the flow from *A* to *B* channeled via an intermediate gene/protein *C*_*i*_. Also, let us denote by 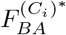 the flow from *A* to *B* channeled via *C*_*i*_, but only through *C*_*i*_-acyclic paths. From the composition properties above, we can write

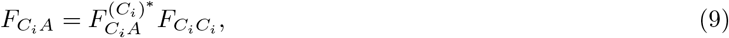

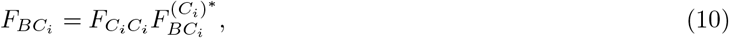

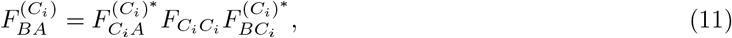

which implies

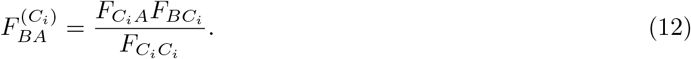

Therefore, the regulation importance 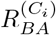 of any intermediate gene/protein *C*_*i*_ in the pathway from *A* to *B*, can be defined as the absolute value of the ratio between the regulation channeled through the intermediate gene/protein and the total regulation,

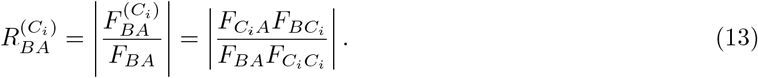

This quantity can be easily calculated, as the terms on the r.h.s. can be read directly from the flow matrix *F* (4).

The relative importance of an intermediate gene/protein allows to design a greedy algorithm to simplify the pathway from *A* to *B* to a collection of successively more precise effective pathways, but comparatively much smaller than the full pathway from *A* to *B*.

### Algorithm 2

Effective pathways between genes/proteins *A* and *B*

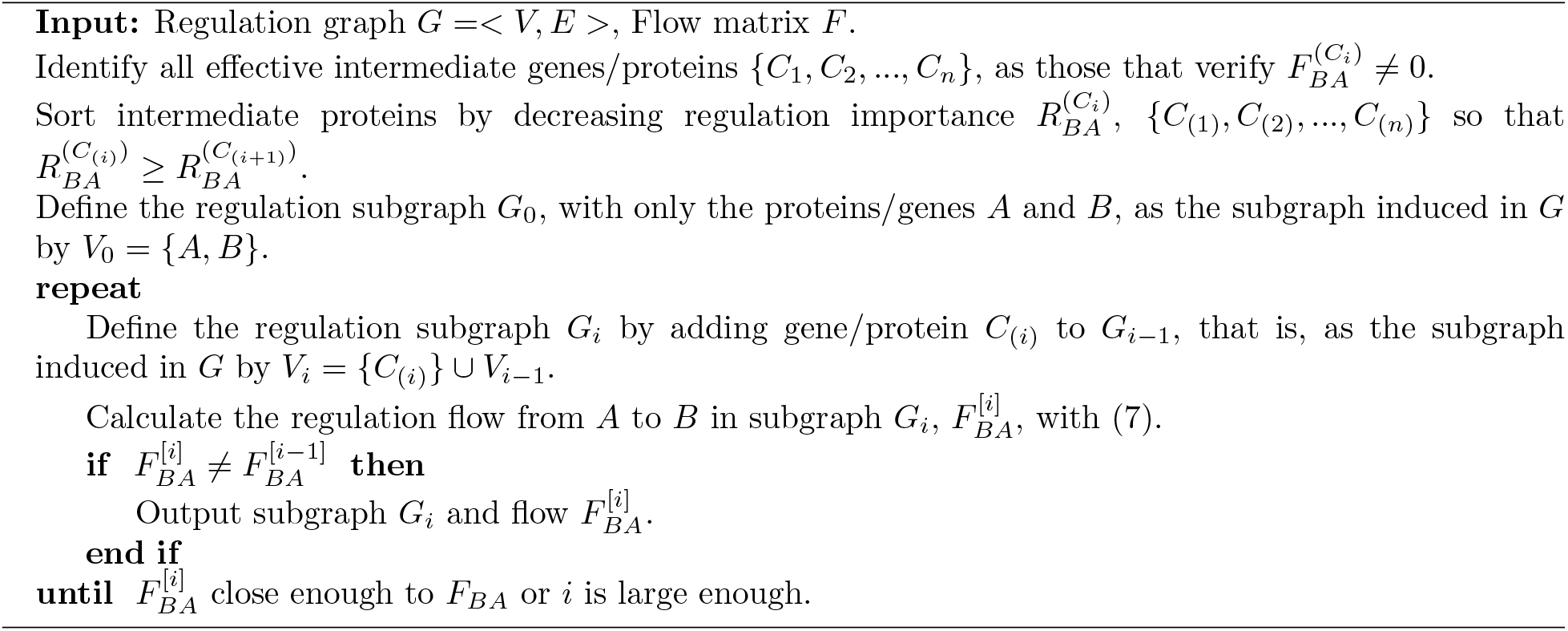

As the edge weights in *G* are signed, some paths may contribute flow which opposes (opposite sign) the overall flow. In those cases the increase in the finite sequence 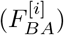 is not monotonous, but convergence is ensured as by construction 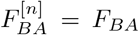. In the initial steps *i* = 0, 1, 2, …, the subgraph *G*_*i*_ may be disconnected, and thus 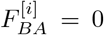. The subgraphs *G*_*i*_ may also contain *loose branches*, that is, series of interconnected genes/proteins *D*_*j*_ not connected in the subgraph to *A* or *B*. Those loose branches are easily identified, as 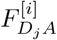 or 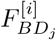 will be zero, and they are no pictured in the generated subgraph.

In summary, after defining the pathway from gene/protein *A* to gene/protein *B* as the subgraph induced by all the intermediate genes/proteins between *A* and *B*, a remarkable capability of graph flow is the calculation of the flow channeled through any intermediate gene/protein. This, in turn, allows to rank intermediate genes/proteins by the amount of regulation channeled through them in the pathway, leading to a greedy algorithm that iteratively builds the much smaller, but functionally equivalent, effective pathways from *A* to *B*.

The definition of a pathway between two genes/proteins as the subgraph with all the regulation paths between those genes/proteins can be generalized naturally to sets of genes or proteins: the pathway from gene/protein set 𝒜 to gene/protein set ℬ is defined as the subgraph formed by all the regulation paths from *A*_*i*_ to *B*_*j*_, for all genes/proteins *A*_*i*_ ∈ 𝒜 and *B*_*j*_ ∈ ℬ. Similarly, the regulation flow from gene/protein set 𝒜 to gene/protein set ℬ can be defined as the regulation flow from all genes/proteins in set 𝒜 to all genes/proteins in set. By denoting the regulation flow from 𝒜 set to set ℬ as *F*_ℬ𝒜_, and given the additivity of graph flow, we have:

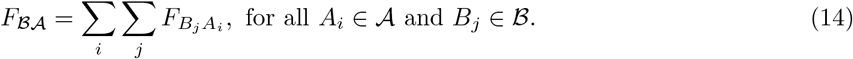

This allows to generalize the Algorithm 2 for identifying effective pathways to sets of genes or proteins, by the corresponding generalization of equation (13): the relative importance of any intermediate gene/protein *C*_*i*_ in the pathway from set 𝒜 to set ℬ, denoted 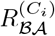, becomes

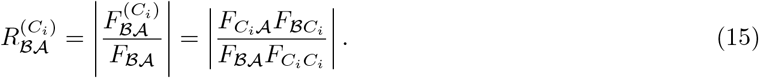

### 3.1 Illustration of the generation of mechanistic hypotheses

To illustrate the algorithm for generating effective pathways, we show how the regulation 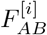 in the successive subgraphs *G*_*i*_ approaches *F*_*AB*_ (see Algorithm 2), in the case of the pathway between TNF and CCL4, i.e. when *A* = TNF and *B* = CCL4.

Figure 2a shows that the regulation fraction 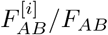 is equal to zero for the initial subgraph *G*_0_ with only the two vertices *A* and *B*, and then it increases non-monotonically to a value close to 1 when the subgraph contains 11 vertices. Thus, this small subgraph explains the regulation flow of the entire subgraph between TNF and CCL4, which has 3,667 vertices, with an accuracy larger than 95%. Figures 2b-2d picture the actual subgraphs constructed by the algorithm at various iterations.

**Figure 1.**
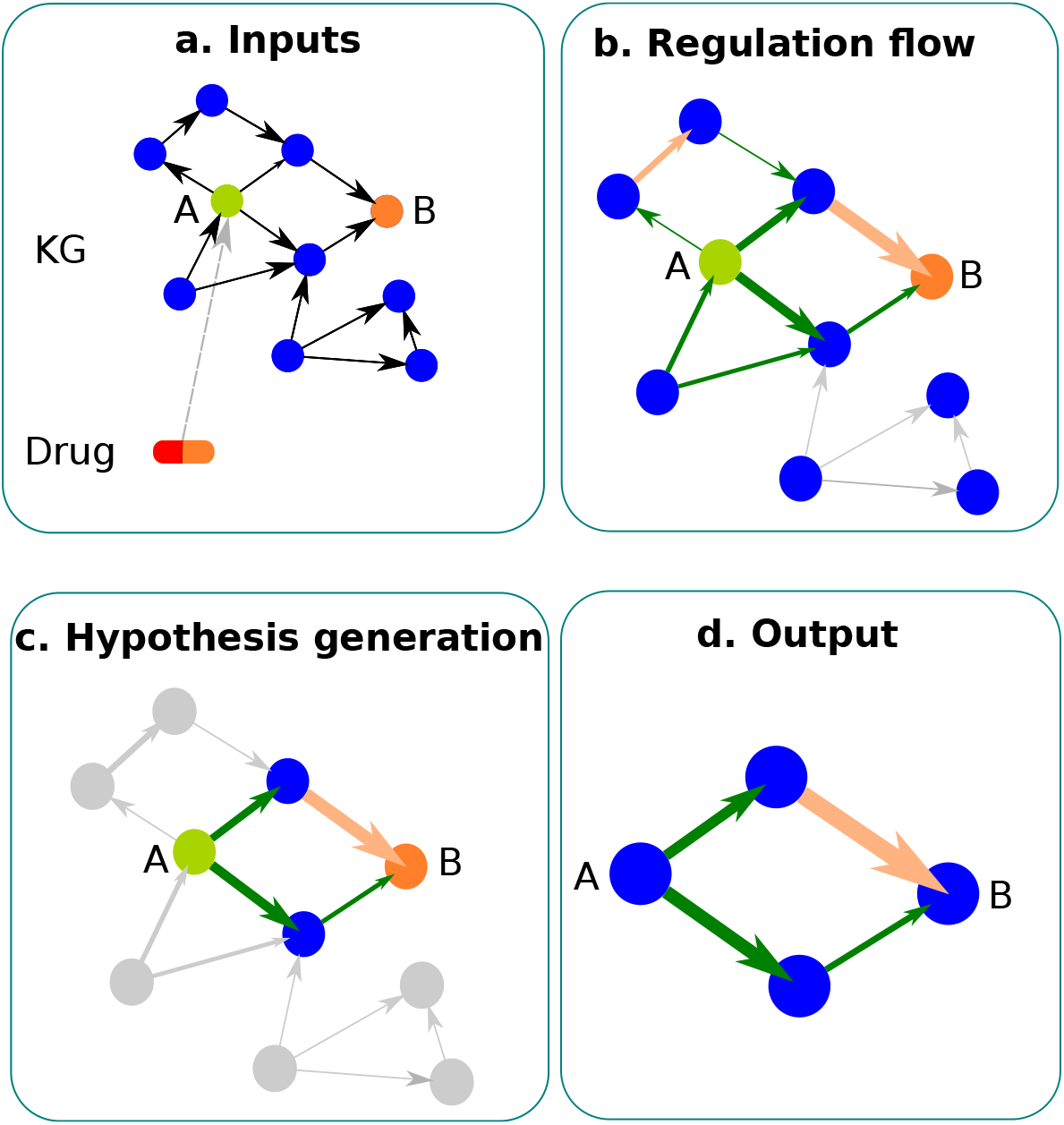
Graphical abstract of the paper: **a**. The Knowledge Graph (KG), the target of the drug (A), and the endpoint of the pathway (B) are provided as inputs; **b**. Regulation Flow is computed in the KG; **c**. Mechanistic hypothesis generation (Algorithm 2) computes the most informative pathways between A and B; **d**. The resulting pathways are output for interpretation and decision making.

**Figure 2.**
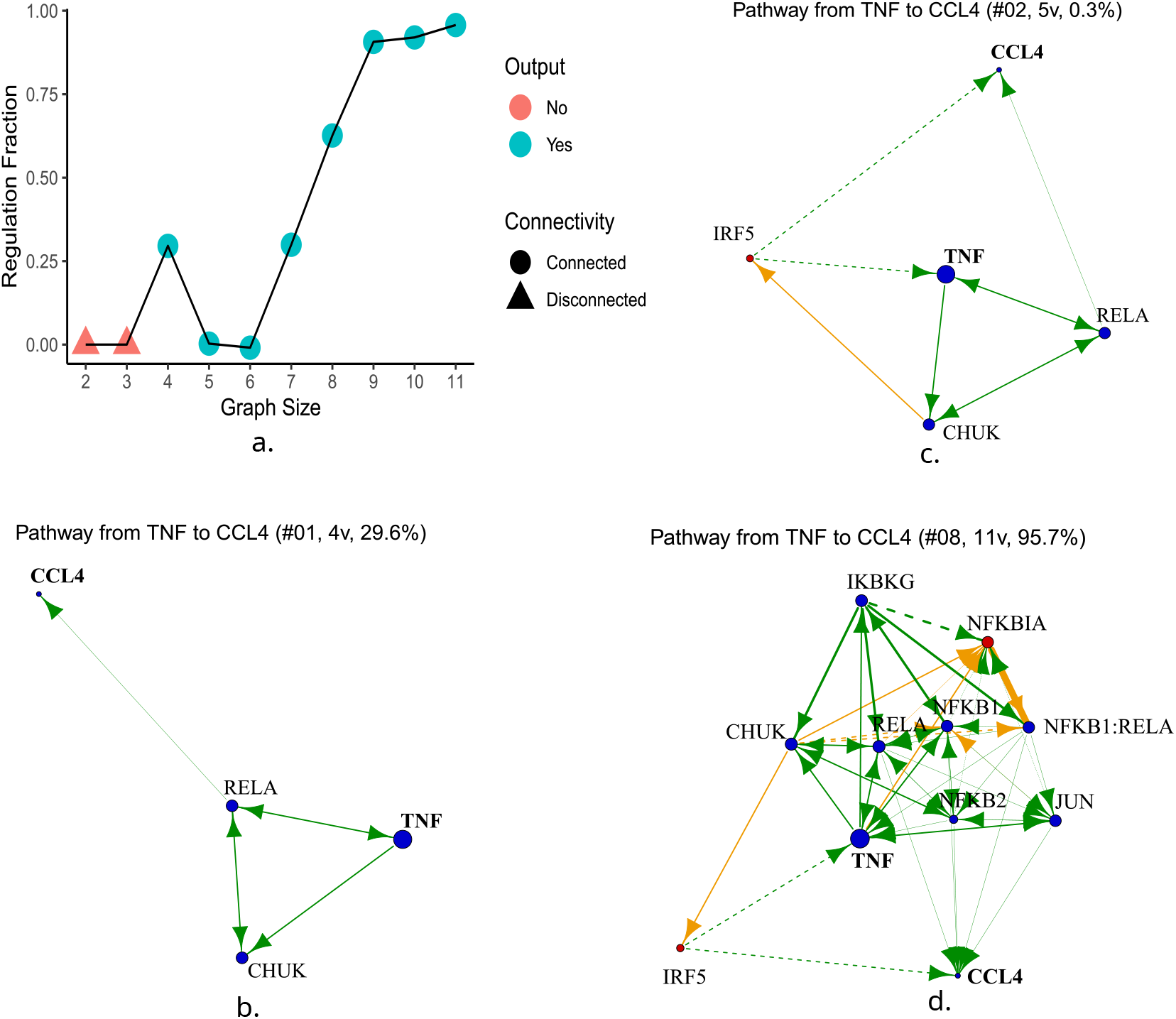
**a**. Dependence of the Regulation Fraction on the subgraph size. When a subgraph is disconnected, it is not output in our implementation. **b**.-**d**. Pathways between TNF and CCL4 with 4, 5 and 11 vertices, respectively. The title is reporting the iteration number (#XX), number of vertices (XXv) and regulation percentage of the given subgraph. Vertices represent genes/proteins, and edges represent direct regulation between genes/proteins. Vertex sizes display the amount of total regulation caused by the original perturbation, TNF. Vertex colors indicate if vertices are regulated in the same way as TNF (blue) or in the opposite way (red), that is, up- or down-regulated, respectively. Edge widths represent the amount or strength of regulation. Edge colors indicate the sign of regulation, positive (green) or negative (orange).

We have observed in our numerical experiments that the regulation fraction changes non-monotonically, and sometimes the regulation fraction can exceed 1. This happens due to the signed nature (stimulation or inhibition) of the biological regulation modeled by the graph: when edges with opposite sign are added in subsequent iterations of the algorithm, newly added paths may channel flow of opposite sign, which counteracts the regulation flow between *A* and *B* so far. This can be observed for example with the subgraphs #01 and #02 in Figs. 2b and 2c.

## 4 Results

Validation of RFA is shown by demonstrating the mechanism of action of genes that have been implicated in causing Noonan Syndrome and related disorders. Syndromes including Noonan Syndrome, Costello syndrome, LEOPARD syndrome and neurofibromatosis type 1 are caused by germline mutations in genes that cause dysregulation of the RAS-MAPK pathway and are collectively know as RASopathies [21]. In addition, an example of applying RFA in a pharmaceutical setting is provided by exploring targets for Fragile X syndrome.

### 4.1 RFA recovers known mechanisms of action of RASopathies

Signaling through the RAS-MAPK pathway plays a critical role in cell differentiation, proliferation and survival. The various mutations in 22 genes that have been associated with RASopathies (Table 1) cause an up-regulation of the RAS-MAP pathway, with *MAPK1* and *MAPK2* being dysregulated downstream. Here we use the known biology of RASopathies to validate RFA, by confirming that the method can reconstitute the expected pathway and predict how the mutations act to cause overall downstream dysregulation of *MAPK1*.

**Table 1:**
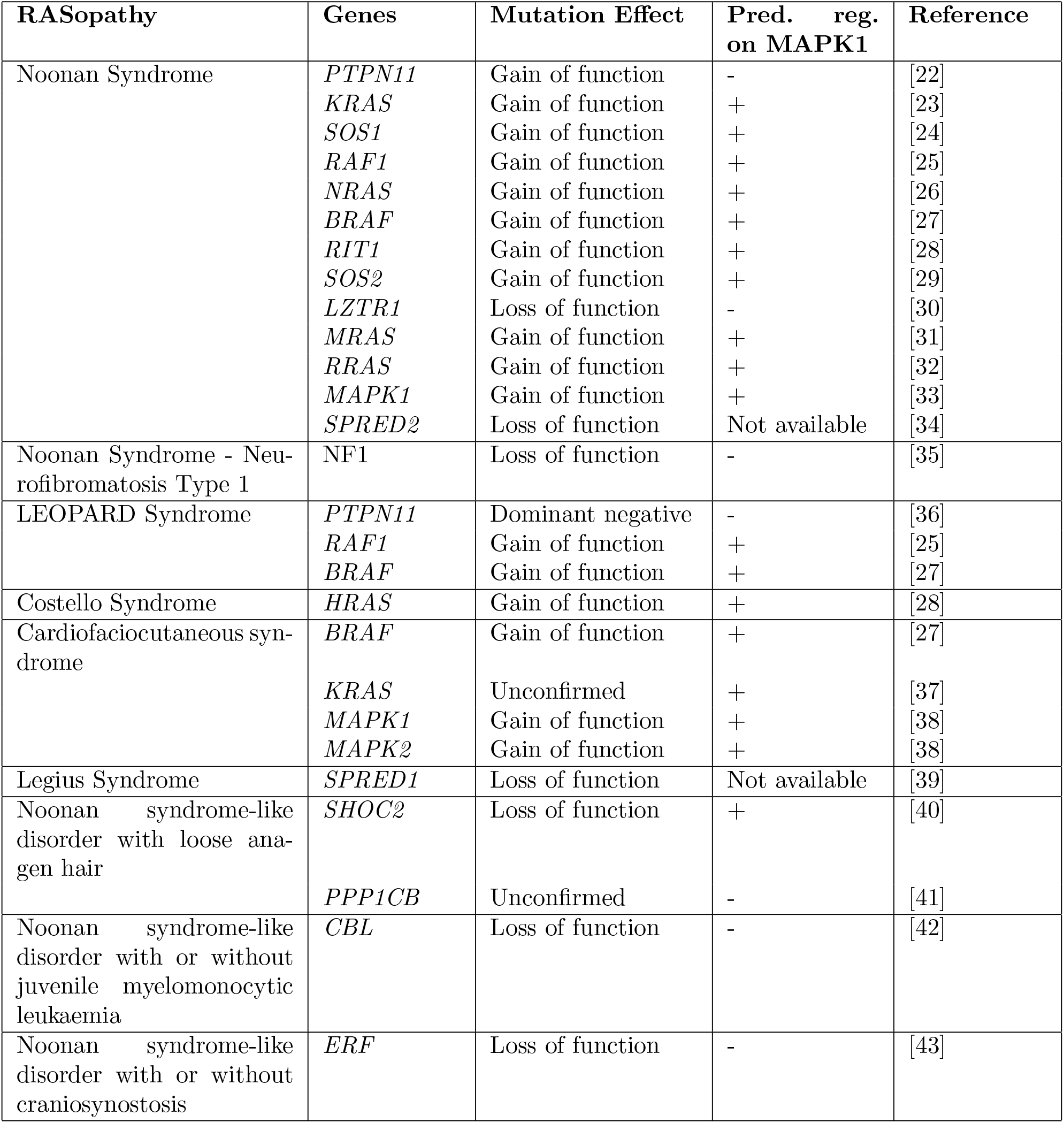
Noonan-like syndromes and the genes that have been implicated in the disease.

The results from RFA are shown in Figure 3. No pathways between *SPRED1* or *SPRED2* to *MAPK1* were generated, as they were not annotated in OmniPath. The results confirm the expected mode of regulation of *MAPK1*, except in the case of two genes; *PTPN11* and *SHOC2*. RFA predicts that overall *PTPN11* has a negative effect on the regulation of MAPK1. The *PTPN11* mutations that cause Noonan syndrome have been shown to be gain of function mutations, which overstimulate the MAPK/ERK pathway. Kontaradis et al. [36] demonstrate that in LEOPARD syndrome the mutations are limited to the PTP domain and have a dominant negative effect by inactivating the catalytic function. This raises the question of how gain of function and dominant negative mutants can result in a similar phenotype. Kontaradis et al. argue that Noonan syndrome and LEOPARD syndrome phenotypes result from differential effects of mutant Shp2 on different receptor-tyrosine kinase pathways at distinct developmental times. The pathway derived by RFA (Figure 3b) indicates two possible routes of regulation between *PTPN11* and *MAPK1*., one of which acts through HRAS and would require a gain of function mutation in *PTPN11*, while the second pathway requires *VAV1*. AS *PTPN11* inhibits the action of *VAV1*, which in turn up-regulates *MAPK1*, a loss of function within *PTPN11* would result in increased activity of both *VAV1* and *MAPK1*. It is interesting to hypothesise that in LEOPARD syndrome it is the *PTPN11* /*VAV1* pathway that is regulating *MAPK1*. RFA predicts that overall *SHOC2* up-regulates *MAPK1*, although Cordeddu describe the mutations as loss of function. The shortest pathway generated by RFA illustrates *SHOC2* acting through *PPP1CA* to down-regulate *MAPK1* (Figure 3c). In this case, the results agree with the literature. The graph shows a dotted yellow line between *PPP1CA* and *MAPK1*, as the predicted inhibition is opposite to the overall effect. Adding extra nodes to the graph shows that there are alternative pathways between *SHOC2* and *MAPK1*, and we hypothesize that the pathway through *PPP1CA* shown in Fig. 3c is the one causing the RASopathy. Functional studies have not been performed on the mutations in *PPP1CB*, whereas RFA results predict that loss-of-function mutations would result in up-regulation of ERK. The recent discovery that loss-of-function mutations in *ERF* cause a Noonan-syndrome-like disorder, with or without craniosynostosis, led Dentici et al. [43] to suggest that this disorder should be classified as a RASopathy. The results shown in Fig. 3a confirm that the mode of action is similar to other RASopathies and support the suggestion by [43].

**Figure 3.**
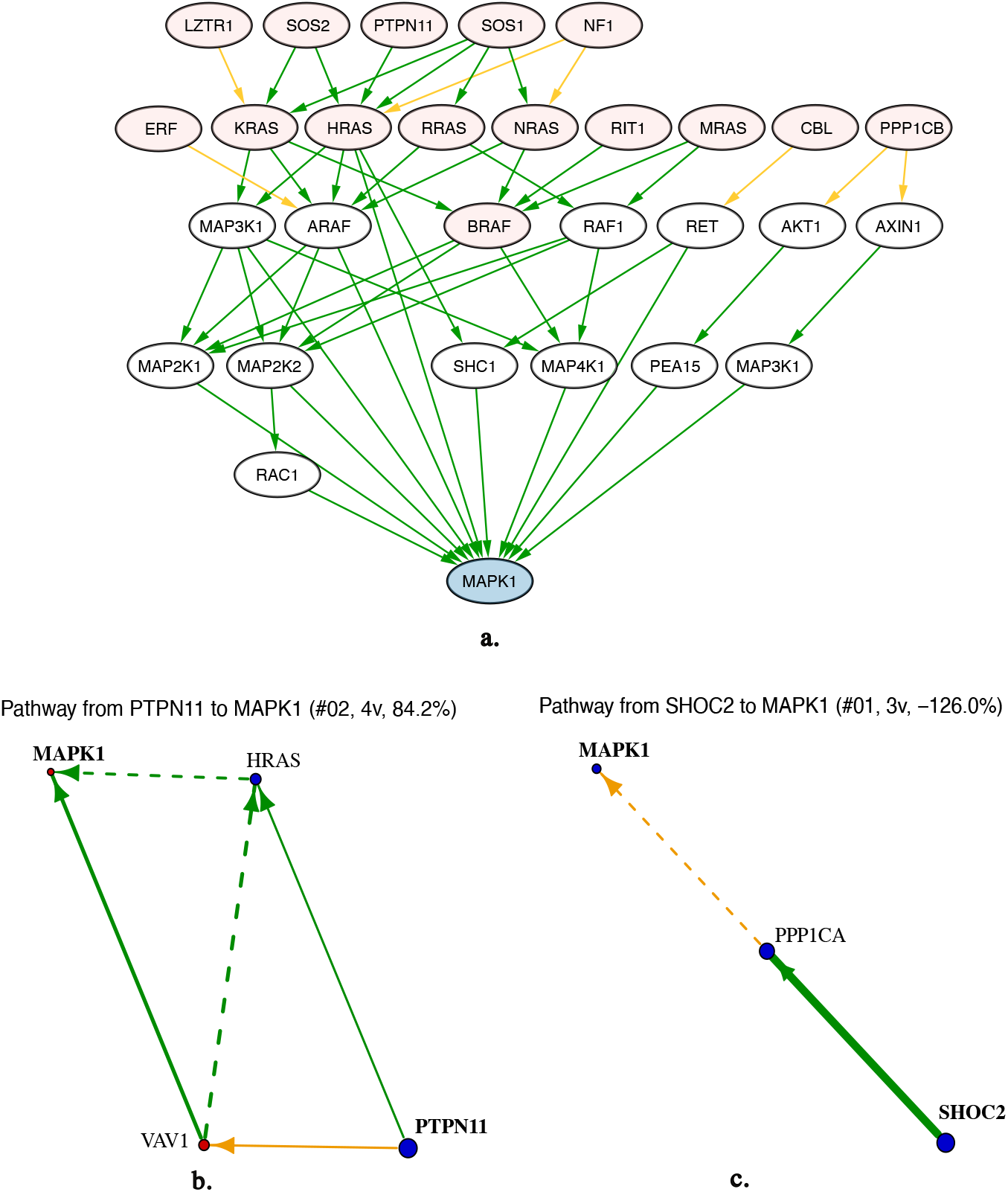
RFA analysis predicts the mutation effect in genes causing RASopathies. **a**. This figure, generated by Cytoscape with RFA results (by integration of the vertices and edges from different RFA results), illustrates interconnected signaling pathways involved in RASopathies, and highlights critical components, including the RAS family to MAPK pathway (pink and blue vertices). Dysregulation of these pathways leads to the pathogenesis of RASopathies, such as Noonan syndrome. Pink vertices in upstream position correspond to genes with known association to RASopathies. We used those genes as source genes in RFA, and MAPK1 as the target gene. Green lines indicate positive regulation, yellow lines indicate negative regulation. Arrows indicate flow direction. **b**. and **c**. show the pathways produced by RFA from PTPN11 and SHOC2, respectively (sizes, widths, and colors of vertices and edges are as defined in Fig. 2)

### 4.2 RFA discovers targets for Fragile X syndrome

Fragile X syndrome (FXS) [44] is caused by mutations in the *FMR1* gene, leading to a non-functional FMRP protein. Gene therapy has been proposed as a way to recover the expression of FMRP. From a drug development and commercial perspective, gene therapy is expensive, and in the case of FXS the therapy would need to be delivered across the blood-brain barrier. We used RFA to investigate whether there were other possible drug target candidates within the regulation pathway downstream of *FMR1*. Brain-derived neurotrophic factor (BDNF) has been proposed as a biomarker for FXS [45], and it has been shown to be dysregulated in *FMR1* knock-out mice [46]. Therefore we investigated how *FMR1* regulates *BDNF* expression, with the aim of uncovering as potential targets the main intermediate genes/proteins channeling this regulation.

Figure 4 shows that *FMR1* up-regulates *BDNF* expression through *PAK1* and *DLG1. DLG1* encodes for a scaffold protein that contains a PDZ domain and is implicated in synaptic activity. Small molecule and peptide inhibitors of PDZ activity have been developed for neurological and cancer indications [47]. *PAK1* encodes for a protein kinase known to play a wide role in cell signaling through its catalytic and scaffolding activities, and it is known to be dysregulated in various neurological disorders including FXS [48]. PAK-inhibiting compounds have been developed for both neurological disorders and cancer [49]. Genentech has developed PAK-inhibitors specifically to treat FXS, which permeate across the blood-brain barrier and ameliorate the FXS phenotype in *Fmr1* knock-out mice [50].

**Figure 4.**
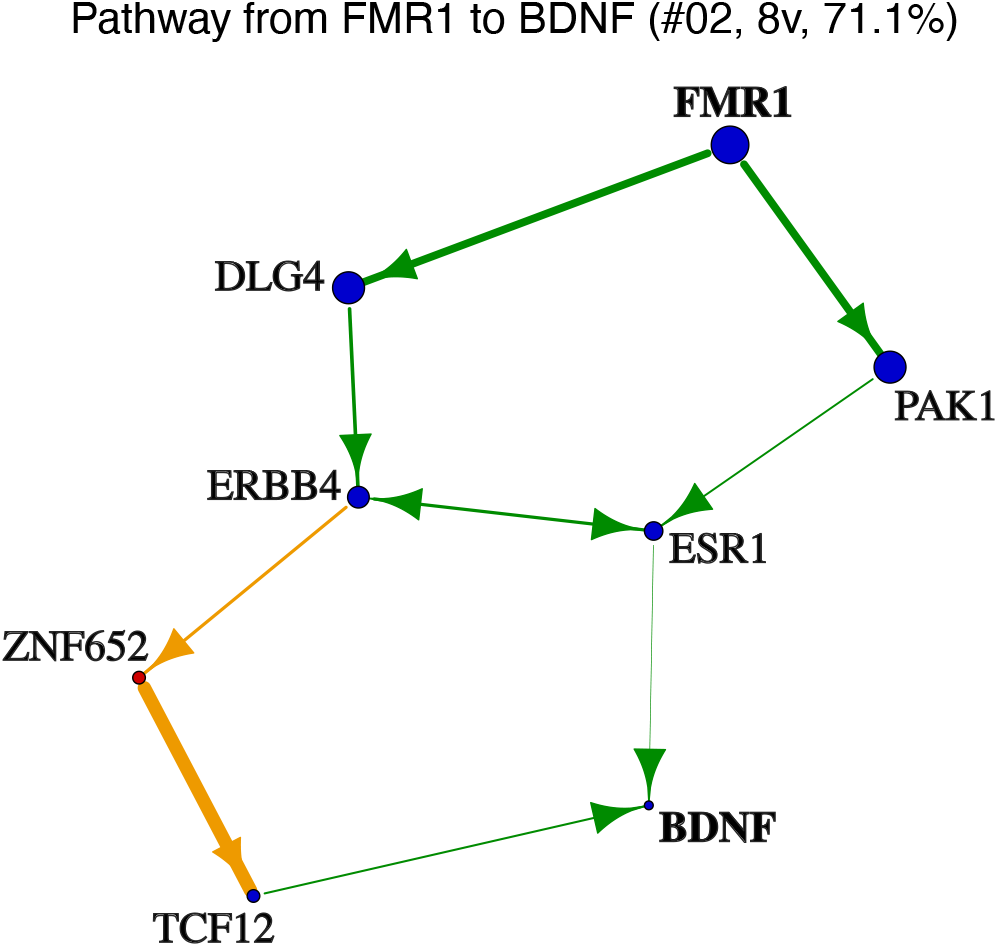
Regulation pathway from *FMR1* to *BDNF*. Sizes, widths, and colors of vertices and edges are as defined in Fig. 2.

In summary, the pathway generated by RFA shows how *FMR1* regulates *BDNF* and allows for the identification of intermediate proteins that have the potential to act as drug targets. Importantly, the causal and molecular nature of the hypotheses generated by RFA allows for clearly defined experimental validation.

## 5 Discussion

We have presented here the Regulation Flow Analysis (RFA) model, which aims to quantify the amount of regulation between two genes or proteins, by making use of regulation graphs and their flow properties. By using the RFA model, we have devised a greedy algorithm for generation of mechanistic hypotheses.

One of the main findings of our paper is that this algorithm is often able to construct *effective pathways* between two biological units of interest, which are much smaller than the full pathway between them, while representing a similar amount of regulation. For example, the full pathway from TNF to CCL4 includes 3,367 genes/proteins, but more that 95% of regulation in this pathway can be explained by a pathway with only 11 genes/proteins. This demonstrates that our algorithm can be seen as a powerful tool for generating understandable and testable models of mechanisms of action.

The application of graph flow carries an implicit assumption of linearity, in the sense that direct regulation between any two connected genes/proteins can be quantified by the multiplication of a factor (the edge weight), and that the regulation over any gene/protein from several direct upstream connections can be calculated by their sum. Apart from the edge weights, RFA has a main parameter, *α* in Eq. (5), which ensures a realistic model with finite total regulation. Qualitatively, this parameter reflects the overall scale of the response produced by perturbations.

Our proposed method, RFA, and the Signal Flow Control (SFC) method published previously [18, 19] have certain commonalities and differences. Based on principles of signal propagation and gradient computations, the SFC authors derive Eq. (7) in [19], which seems similar to our Eq. (6). However, the modeling approaches of RFA and SFC are different. On one hand, RFA aims to model biological regulation of any kind, not only signaling processes. On the other hand, RFA models the effect of a given perturbation simply as the propagation of regulation, whereas SFC models this effect as a user-defined combination (sum) of the propagation of regulation and basal activity. As a consequence, SFC has two user-defined hyperparameters (*α* and *β* Eq. (7) in [19]) defining that combination, while RFA has none of this. The authors of SFC note that their flow computation is not possible directly sometimes. Indeed, in some cases the user-defined hyperparameters may lead to the scaled step matrix having a spectral radius larger than 1, and therefore its inverse will not exist. Possibly due to this, SFC authors propose an iterative algorithm involving a repeated matrix multiplication (Table 3 of [19]), rather than the direct inverse computation carried out in RFA. Depending on the amount of iterations, this may lead to a much greater computational cost compared to the direct inversion, and it also may lead to numerical overflow if the spectral radius of the scaled step matrix is close to 1. Another, less important, difference between RFA and SFC is the different scaling of the step matrix, using only out-degree (RFA) or both in-degree and out-degree (SFC).

We have performed some comparison of performance with real datasets, using the default hyperparameters values in SFC, which emphasize regulation flow over basal activation. As expected with that setting, results show that the quality of prediction of RFA and SFC, in terms of sign and magnitude of perturbation effects over a large set of genes, are similar: one model performed better in some scenarios, while the other model had better performance in others, resulting in similar overall performance of the two methods (see Supplementary Information).

The two provided examples about RASopathies and Fragile X Syndrome showcase the potential of RFA in drug research and development. The molecular nature of regulation graphs, together with their directionality and signed nature, establish a general framework for understanding the causal relationships between genetic variation, disease mechanisms, and biomarkers. As a consequence, it facilitates the rational discovery and validation of disease targets, in the form of genes/proteins predicted to have a therapeutically beneficial regulation influence on the core events driving a disease. Compared to Bayesian graphs, where edge weights represent conditional probabilities, regulation graphs provide a more adequate framework to model biological regulation, given the built-in flexibility in the sign and weight of edges.

RFA, like other models based on regulation graphs, is highly dependent on a good estimation of the edge weights, that is, on the proper estimation of the strength of regulation between genes/proteins. This problem can be addressed by refining the estimation using machine learning and artificial intelligence approaches, leveraging as training data publicly available experimental libraries, with gene expression profiling of cell lines stimulated with a variety of factors [51]. This way, the compact matrix algebra of RFA would enable to train causal AI models, with direct molecular interpretation and well-defined empirical validation. Other possible directions of exploration are generalizations of the RFA model to take into account the distinction between inter- and intra-cellular regulation, as well as the differentiation between cell types.

Thus, we envision that RFA-based approaches will become prevalent and more sophisticated, in the pursuit of deeper understanding and more precise modeling of disease biology at molecular and cellular level. These efforts, hand in hand with extensive omics profiling, will facilitate rational drug design, with substantial improvements in the duration and success rates of drug projects.

## 6 Supplementary information

To compare prediction accuracy between the RFA and SFC algorithms, we used a dataset introduced in [52]. This dataset contains 66 different perturbations of a small network with 22 vertices, with each perturbation being performed under 200 secondary conditions, corresponding to different dose levels of EEGF and insulin, and different exposure times. Accordingly, the dataset contains measurements of 66*×*200 = 13, 200 perturbation experiments. In each of those, we obtained the Regulation Flow with RFA and the Direction of Activity Change with SFC. The predicted values from both algorithms were used to calculate: a) mean accuracy, by comparing the sign of the prediction with the sign of the changes observed experimentally in genes/proteins (graph vectices); b) Spearman’s correlation for the ordinal association between predicted and observed changes in genes/proteins. The results presented in Figure 5 illustrate that RFA and SFC have similar performance based on both metrics.

**Figure 5.**
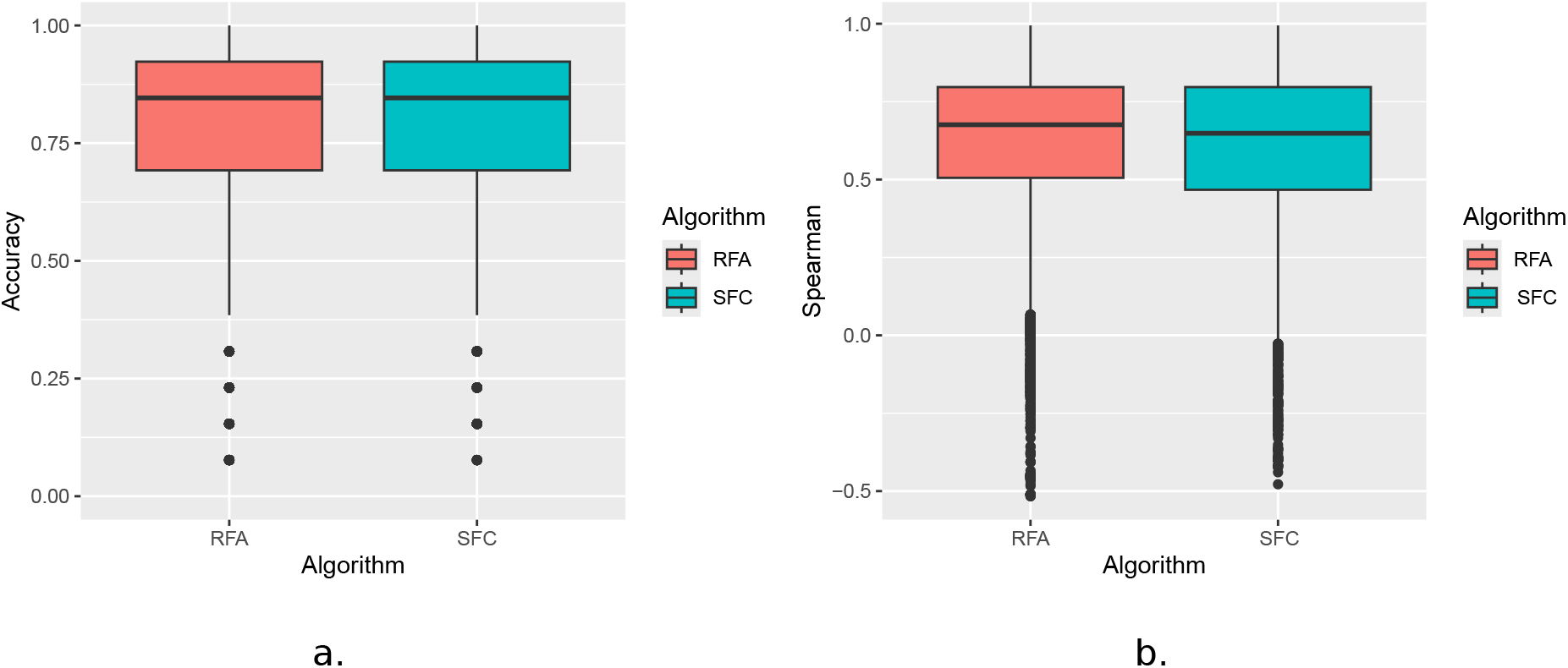
Comparison of RFA and SFC methods in terms of: **a**. accuracies of sign of predicted expression changes, and **b**. Spearman’s correlation between observed and predicted expression changes.

## 7 Code availability

An implementation of RFA is available at https://github.com/AstraZeneca/regflow.

